# *Russula* subgen. *Cremeoochraceae* subgen. nov.: a very small and ancient lineage sharing with *Multifurca* (*Russulaceae*) an identical, largely circum-Pacific distribution pattern

**DOI:** 10.1101/2023.07.26.550279

**Authors:** B Buyck, E. Horak, J.A. Cooper, X.H Wang

## Abstract

The application of DNA data on a worldwide sampling has revolutionized the infrageneric classification of the highly diverse ectomycorrhizal genus *Russula*. Based on collections made in New Zealand, East Asia and North America, this study describes the new subgenus *Cremeoochraceae*, the ninth subgenus in *Russula*. Even though BLASTn of the ITS sequences suggested affinities with species of subgenera *Russula* and *Heterophyllidiae*, the phylogenetic analysis based on a five-locus DNA dataset placed the target samples in an independent major clade that is systematically equivalent to subgenus. The new subgenus shares with subgen. *Brevipedum* subsect. *Pallidosporinae* the general field habit, the unequal lamellae and the relatively small spores with inamyloid suprahilar spot and similar spore ornamentation. It differs from the latter subsection principally in the poor contents of all types of cystidioid cells and the often areolate-scurfy pileus surface composed of slender, undulating hyphal terminations with frequent subcapitate apices. Biogeographically, subgen. *Cremeoochraceae* shares with *Multifurca*, the sister genus of *Lactarius*, a circum-Pacific distribution pattern with the exception of South America. Both lineages lack representatives in Europe and Africa. The hypothesis proposing an African origin for the genus is considered unlikely.

## Introduction

Once the question of the monophyly of *Russula* had been settled (Buyck *et al*. 2008), the traditional and highly artificial infrageneric classifications of the genus (e.g. Singer, 1932, 1986; Romagnesi 1967, 1987; Sarnari 1998, 2005) have been rapidly abandoned for a more natural phylogeny based on multilocus DNA data (Bazzicalupo *et al*. 2017; Looney et al., 2016; Buyck *et al*. 2018, 2020). Molecular techniques, however, were not the only aspect that contributed to a more natural classification. A representative sampling of the world’s diversity of the genus was an equally important aspect toward the recognition and documentation of the higher infrageneric taxa (Wang *et al*. 2019; Rossi *et al*. 2020).

The classification of *Russula*, one of the most diverse fungal genera (He *et al*. 2019), will certainly continue to benefit from the ongoing description and sequencing of new and rare or ill-known species. Already in 2020, the discovery of a Chinese look-alike of the very rare North American *R. glutinosa* ultimately resulted in the description of the new subgenus *Glutinosae*, an ancient lineage comprising at present merely two species (Buyck *et al*. 2020).

It is again the rediscovery of another very rare North American *Russula* that resulted in the present contribution. It all started with the publication on the website of iNaturalist of an unidentified *Russula* collected in New York State (https://www.inaturalist.org/observations/56944584). Having received part of this collection, the first author (BB) could rapidly identify the specimen as being very close or identical to *R. inopina* Shaffer thanks to a recently published type-study of that species (Buyck & Adamcik 2013). *Russula inopina* was originally considered a close relative to *R. brevipes* and was never recollected since its original description by Shaffer (1964). In the field and under the microscope, Shaffer’s species is indeed reminiscent of species belonging to subgen. *Brevipedum* subsect. *Pallidosporinae*, but it is the unexpected BLASTn result of the ITS sequence that triggered our interest in this collection. This paper addresses the systematic placement of this *R. cf. inopina* and its closest relatives and discusses its impact on the biogeographic history of the genus.

## Material and Methods

### Morphology

Microscopic characters were examined under a Nikon E400 microscope (Nikon, Tokyo) at a magnification of 1000 ×. Fragments of hymenium and pileipellis were shortly heated in a 5% KOH solution before observation in Congo red ammonium solution. Spore ornamentation was observed in Melzer’s reagent. Line drawings were made at a projected magnification of 2400 × with the aid of a drawing tube (Y-IDT, Nikon, Tokyo, Japan). Spore dimensions are given following the form (a) b−***m***−c (d), with ***m*** the mean value, b−c containing at least 90% of all values and with extreme values (a, d) enclosed in parentheses. Q indicates the basidiospore length/width ratio.

### Sampling

For a representative sampling of all known subgenera of *Russula*, we included sequences of 85 species. Among these, sequence data of six species were retrieved from the database of JGI (https://genome.jgi.doe.gov/portal/). These species were preliminary identified as *R. brevipes* (BPL707), *R. compacta* (BPL669), *R. dissimulans* (BPL704), *R. earlei* (BPL698), *R. seminuda* (PSC4341-TAS1) and *R. vinacea* (BPL710). The identifications of the abovementioned American species are to be interpreted ‘sensu lato’ as these names still correspond to species complexes. Sequences of the other specimens were borrowed from Buyck *et al*. (2018), Wang *et al*. (2019), Buyck *et al*. (2020), Rossi *et al*. (2020) and Cao *et al*. (2023) or newly generated in this study. To guarantee more solid evidences for phylogenetic inference, we specifically selected samples that could minimize missing data in our multigene phylogeny. Considering the limited sampling for *R*. subgen. *Crassotunicatae* in Buyck *et al*. (2018, the five genes were only available for *R. farinipes*), we sequenced two Chinese samples (XHW8474 and XHW4498) that are closely related to *R. pallescens* and *R. crassotunicata*, respectively. We also added multi-gene data for *R. maguanensis, R. substriata* and *R. shingbaensis*, three Asian members of subsection *Substriatinae* of subgen. *Heterophyllidiae*, as well as for the Chinese *R. ochrobrunnea* of sect. *Polyphyllae* of subgen. *Compactae*. New sequences were also generated for several species in subgen. *Brevipedum* (*R. flavescens, R. pseudojaponica, R. brevispora* and *R*. aff. *pallidospora*) because of the shared morphology between this subgenus and *R. cf. inopina*. Because of the high similarity in ITS sequences and the similar micromorphology shared between *R. cf. inopina* and *R. cremeoochracea* from New Zealand, we sequenced three specimens of the latter species. One Chinese sample (QC376) that was molecularly similar was added to the dataset. New sequences generated in this study are listed in Table I.

### DNA extraction, PCR amplifications, and sequencing

For DNA extraction, PCR and sequencing, we basically followed the methods and primers of Buyck *et al*. (2018). For problematic samples, internal primers designed by Cao *et al*. (2023) were used to separately amplify shorter sections within each locus. We amplified five loci to infer the phylogenetic relationships among the species, following Buyck *et al*. (2018, 2020): mitochondrial rDNA small subunit (mitSSU), nuclear rDNA large subunit (nucLSU), RNA polymerase II largest (*rpb*1) and second largest subunit (*rpb*2), and translation elongation factor 1-alpha (*tef1*). In addition, we amplified ITS of three samples of *R. cremeoochracea*, one specimen corresponding to *R. cf. inopina* and one specimen of an undescribed species (Chinese sample QC376) to aid in phylogenetic species recognition.

PCR amplification and Sanger sequencing for the three specimens of *R. cremeoochracea* (ZT 68-038, ZT 69-002 and ZT 69-36) and the holotype of *R. ochrobrunnea* (GDGM 79718) failed or partly failed. For these specimens, lower coverage whole genome sequencing (“genome skimming”) using the next-generation sequencing by Illumina NovoSeq-5500 platform was used to obtain the missing genes. Library preparation followed the protocol described by Zeng *et al*. (2018). The raw data were de novo assembled using GetOrganelle toolkit (Jin et al., 2020). The target regions were extracted using the reference data from GenBank (MW683813, MW683817, MW683650 and MW683654) and those of *R. cf. inopina* from MF84640.

### Phylogenetic analyses

Sequences generated by Sanger sequencing were assembled and edited using Sequencher (Gene Codes Corporation, Ann Arbor, USA). Alignments were performed using ClustalW in BioEdit (Hall 1999) and manually adjusted. Two datasets were used: an ITS-nucLSU dataset to aid species delimitation within terminal clades and a five-locus combined dataset (nucLSU, mitSSU, *rpb1, rpb2* and *tef1*) for phylogenetic inference at genus level. The ITS-nucLSU alignment was partitioned into ITS and nucLSU. For the five-locus dataset, we followed Buyck *et al*. (2018) to exclude ambiguous sites and introns and to set the best partitioning, i.e. nucLSU, mitSSU, *rpb1* intron, *rpb1* 1st, *rpb1* 2nd, *rpb1* 3rd, *rpb2* 1st, *rpb1* 2nd, *rpb1* 3rd, *tef1* 1st, *tef1* 2nd and *tef1* 3rd codon positions. Phylogenetic trees were rooted in the ITS-nucLSU dataset using two samples of *R. glutinosoides* and, in the five-locus dataset, with three species of *Lactifluus*. All best tree searches and bootstrap analyses were conducted in RAxML version 7.7.7 (Stamatakis *et al*. 2008) using the rapid bootstrap algorithm (RBS). The general time-reversible substitution model with site rate heterogeneity was selected (option -m, GTRGAMMA) and 1000 runs with distinct heuristic starting trees (option -n 1000) were executed. Combinability among the five datasets was examined conducting 1000 RBS heuristics for each single locus dataset. Conflict was assumed when single-locus genealogies for the same set of taxa inferred different relationships (monophyletic versus non-monophyletic) both with significant RBS support values (RBS ≥ 70%; Mason-Gamer and Kellog, 1996). Tree branches were considered significantly supported when RBS values were ≥ 70% (Alfaro *et al*. 2003).

## Results

### Newly produced sequence data

A total of 94 new sequences were generated in this study (5 ITS, 17 nucLSU, 19 mitSSU, 19 *rpb1*, 17 *rpb2* and 17 *tef1*) and deposited in GenBank (see Table 1). These newly produced sequences belong to taxa placed in five known subgenera and the here newly described subgenus.

**Table 1.** Voucher table for newly sequenced specimens used in the multilocus (Fig. 1) and combined ITS-nucLSU phylogeny (Fig. 2). The sequences in bold are generated by this study and the specimens marked with asterisks are sequenced by next generation sequencing.

| Species | Geographical origin | Collection nr. | ITS | nucLSU | mitSSU | *rpb1* | *rpb2* | *tef1* |
| --- | --- | --- | --- | --- | --- | --- | --- | --- |
| *R. brevispora* | Guizhou, Yunnan | X.H. Wang 10183 (KUN-HKAS 122203) | – | **OQ913517** | **OQ916941** | **OQ919750** | **OQ919749** | **OQ919748** |
| *R. cyanoxantha* | Yunnan, China | X.H. Wang 4603 (KUN-HKAS 104888) | – | **OQ359146** | **OQ371392** | **OQ359994** | **OQ360017** | **OQ360007** |
| *R. compactoides* | Yunnan, China | X.H. Wang 5845 (KUN-HKAS 116518) | – | **OQ359144** | **OQ371390** | **OQ359993** | **OQ360016** | **OQ360005** |
| *R. aff. crassotunicata* | Yunnan, China | X.H. Wang 4498 (KUN-HKAS 104770) | – | **OQ359150** | **OQ371396** | **OQ359998** | **OQ360020** | **OQ360010** |
| *R. cremeo-ochracea\** | New Zealand | E. Horak (ZT 69-365) | **OQ371409** | **OQ371409** | **OQ371402** | **OQ359985** | **OQ359989** | **OQ359982** |
| *R. cremeo-ochracea\** | New Zealand | E. Horak (ZT 68-038) | **OQ371410** | **OQ371410** | **OQ371403** | **OQ359986** | **OQ359990** | **OQ359983** |
| *R. cremeo-ochracea\** | New Zealand | E. Horak (ZT 69-002) | **OQ371411** | **OQ371411** | **OQ371404** | **OQ359987** | **OQ359991** | **OQ359984** |
| *R. maguanensis* | Yunnan, China | X.H. Wang 4765 (KUN-HKAS 102277) | – | MH714537 | **OQ586179** | **OQ579058** | MH939989 | MH939983 |
| *R. inopina* | New York, USA | Paula deSanto (MF 84640, CUP 070958) | **OQ357663** | **OQ359155** | **OQ371401** | **OQ360003** | **OQ360024** | **OQ360014** |
| *R. flavescens* | Sichuan, China | Y. Li MYJY20220811-01-03 (KUN-HKAS 128519) | – | **OQ860082** | **OQ863199** | **OQ919746** | **OQ919743** | **OQ919740** |
| *R. ochrobrunnea\** | Guangdong, China | S.Y. Zhou K19071502 (GDGM 79718) | – | **OP723360** | **OQ371405** | **OQ359988** | **OP846962** | **OP723479** |
| *R. aff. pallescens* | Yunnan, China | X.H. Wang 8474 (KUN-HKAS 118094) | – | **OQ359151** | **OQ371397** | **OQ359999** | **OQ360021** | **OQ360011** |
| *R. aff. pallidospora* | Yunnan, China | X.H. Wang 8793 (KUN-HKAS 118398) | – | **OQ860083** | **OQ863200** | **OQ919747** | **OQ919744** | **OQ919741** |
| *R. pseudocompacta* | Yunnan, China | X.H. Wang 4663 (KUN-HKAS 104878) | – | **OQ359143** | **OQ371389** | **OQ359992** | **OQ360015** | **OQ360004** |
| *R. pseudojaponica* | Yunnan, China | G. Wu s.n. (KUN-HKAS 124474) | – | **OQ860081** | **OQ863198** | **OQ919745** | **OQ919742** | **OQ919739** |
| *R. shingbaensis* | Yunnan, China | X.H. Wang 9755 (KUN-HKAS 119312) | – | **OQ359149** | **OQ371395** | **OQ359997** | **OQ360019** | **OQ360009** |
| *Russula sp.* | Yunnan, China | Q. Cai 376 (KUN-HKAS 67940) | **OQ357662** | **OQ359154** | **OQ371400** | **OQ360002** | **OQ360023** | **OQ360013** |
| *R. subpunctipes* | Yunnan, China | X.H. Wang 5017 (KUN-HKAS 111469) | – | **OQ359147** | **OQ371393** | **OQ359995** | **OQ360018** | **OQ360008** |
| *R. substriata* | Yunnan, China | X.H. Wang 4766 (KUN-HKAS 102278) | – | MH714540 | **OQ371394** | **OQ359996** | MH939992 | MH939986 |

#### Phylogeny

##### Five-locus combined phylogeny

The combined five-locus dataset consisted of 87 ingroup samples representing 85 species and three outgroup species of *Lactifluus*. After exclusion of ambiguously aligned regions and all introns, the alignment was 3876 bp long. We did not find supported conflicts among the five single locus genealogies. The analysis yielded a best tree that retrieved the monophyly of *Russula* and each of the known eight subgenera, as well as the here newly introduced subgenus *Cremeoochraceae*, all with full support (ML-BS=100%) (Fig. 1).

**Fig. 1.**
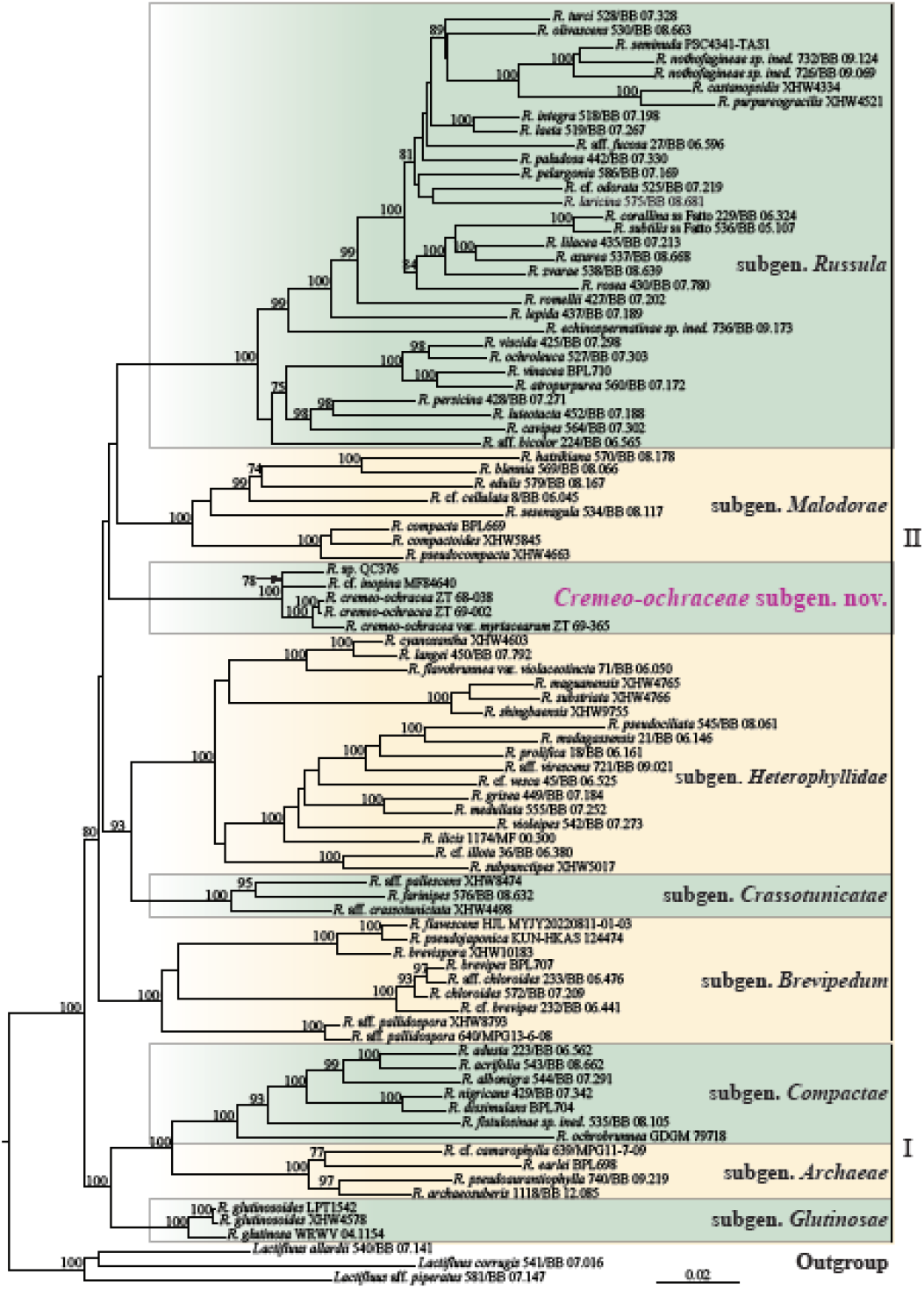
The most likely tree obtained by Maximum Likelihood analysis of the nucLSU-*tef1-rpb2-rpb1*-mtSSU combined dataset, rooted with three species of *Lactifluus*, the sister genus to *Russula*. ML Bootstrap values ≥ 70% are indicated on or by the branches. Infrageneric classification followed Buyck *et al* (2018, 2020).

Our phylogeny divided the genus in two superclades. Superclade I was formed by subgenera *Glutinosae, Compactae* and *Archaeae* and received full ML bootstrap support (ML-BS=100%). Superclade II received moderate, yet significant support (ML-BS=80%) and was composed of our new subgenus and all remaining subgenera (*Russula, Malodorae, Heterophyllidiae, Crassotunicatae* and *Brevipedum*). The short branch (branch length = 0.0036) leading to superclade II was counterbalanced by the strong support obtained for superclade I. Within superclade II, we recovered high support (ML-BS=93%) for the monophyly of *Heterophyllidiae* +*Crassotunicatae* lineage, but the relationships between the other subgenera remained unresolved. The branch length leading to the new subgenus (0.0423) is the longest among those leading to all the individual subgenera.

##### Two-locus, ribosomal gene (ITS-nucLSU) phylogeny

In the combined ITS-nucLSU tree (Fig. 2), one of our samples of *R. cremeoochracea* (ZT 69-365) grouped with significant support (ML-BS = 100%) with two other New Zealand samples (JAC13166 and JAC13170), while the remaining two other samples of *R. cremeoochracea* (ZT 68-038 and ZT 69-002) formed a separate terminal clade (ML-BP=100%), sister to this clade (ML-BP 100%). The two clades differ by five consistent base pairs in the ITS region and four in the LSU region. The American sample of *R. cf. inopina* MF84640 formed a moderately supported clade (ML-BS=71%) with an environmental sequence from Japan (GenBank accession LC315898) and our Chinese unnamed sample QC376. The Malaysian environmental sample with GenBank accession GQ268654 formed a singleton, with unresolved relationships with the other samples.

**Fig. 2.**
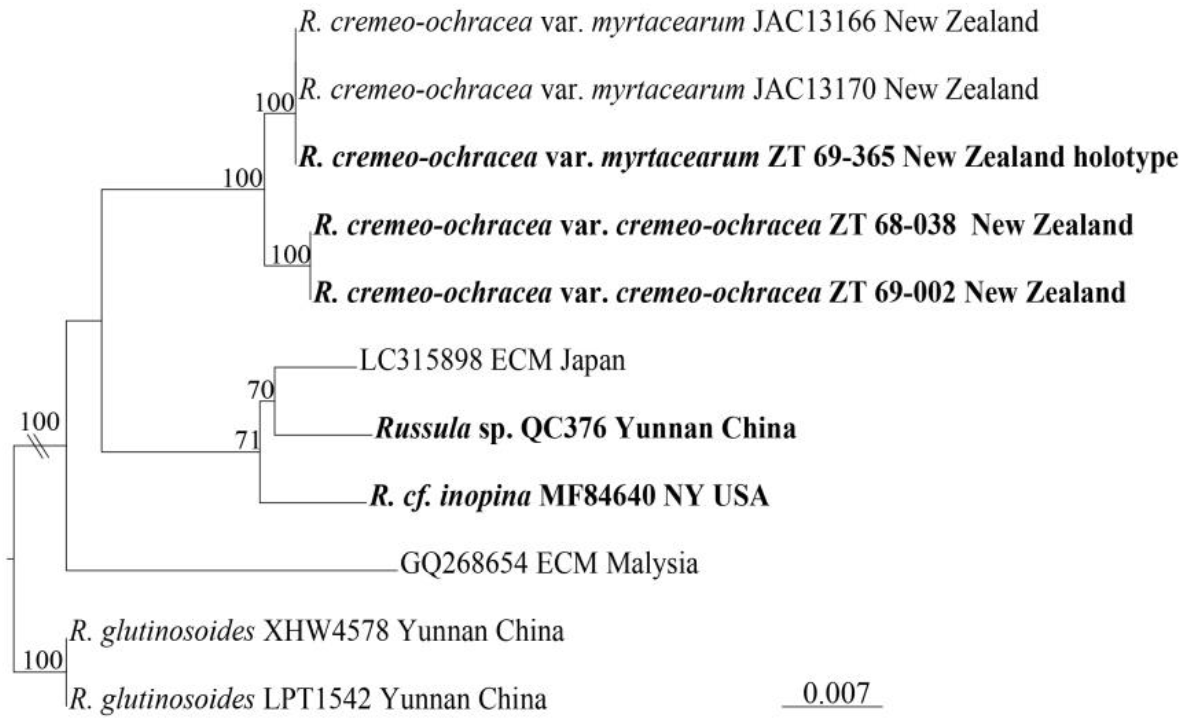
The best tree obtained by the Maximum Likelihood analysis of the combined ITS-nucLSU dataset for members of the new subgenus, rooted with two samples of *Russula glutinosoides*. For the Malaysian and Japanese ectomycorrhizal samples, only ITS sequences are available. Bootstrap values ≥ 70% are indicated on the branches. Specimens sequenced in this study are in bold.

#### Taxonomy

Following the implications of the above phylogenetic analyses (Figs. 1–2), we here describe a new subgenus to accommodate *R. cremeoochracea, R*. cf. *inopina*, and the three still undescribed Asian species.

### *Russula* subgenus *Cremeoochraceae* Buyck & X.H. Wang, subgen. nov. MycoBank: MB xxxxxx

#### Diagnosis

The new subgenus shares with subgen. *Brevipedum* subsect. *Pallidosporinae* the pale-coloured, firm and fleshy basidiocarps with short stipe, the unequal lamellae, the relatively small spores with inamyloid suprahilar spot having a low, subreticulate ornamentation, and the production of a coloured spore print. The new subgenus differs from subsect. *Pallidosporinae* principally in the sometimes poor contents of all types of cystidioid cells and the often areolate-scurfy pileus surface composed of slender, undulating hyphal terminations with frequent subcapitate apices. All of the collected specimens so far have a mild taste.

#### Type species

*Russula cremeoochracea* R.F.R. McNabb, *New Zealand J. Bot*. 11: 683 (1973)

##### Attributed species

### 1. *Russula* cf. *inopina* Shaffer, *Mycologia* 56 (2): 208 (1964). Figs. 3–4

#### Description

**Pileus** medium-sized, fleshy, firm, strongly but narrowly depressed in the center, margin smooth, inrolled when young, then bending downward, never really uplifted; surface dull, smooth or, more frequently, distinctly scurfy-areolate or felty outside the center, particularly so closer to the margin, pale yellow or locally more ochraceous. **Lamellae** adnate, crowded, unequal and polydymous, with 1−3 (7) lamellulae+lamellae/cm, some forking close to the stipe, at first whitish, then turning to a soft ochre yellow (slightly paler to concolorous as compared to the pileus); edges concolorous, entire. Stipe shorter than the pileus diameter, thick and fleshy, cylindrical, white, in the lower part with faint yellowish stains as on pileus, longitudinally rugulose, broadly rounded at the base. **Context** white, firm, unchanging or slightly browning in the insect-damaged areas. **Odor** not remarkable. **Taste** mild. **Spore print** not obtained, but not white (as deduced from the colour change of the gills).

**Fig. 3.**
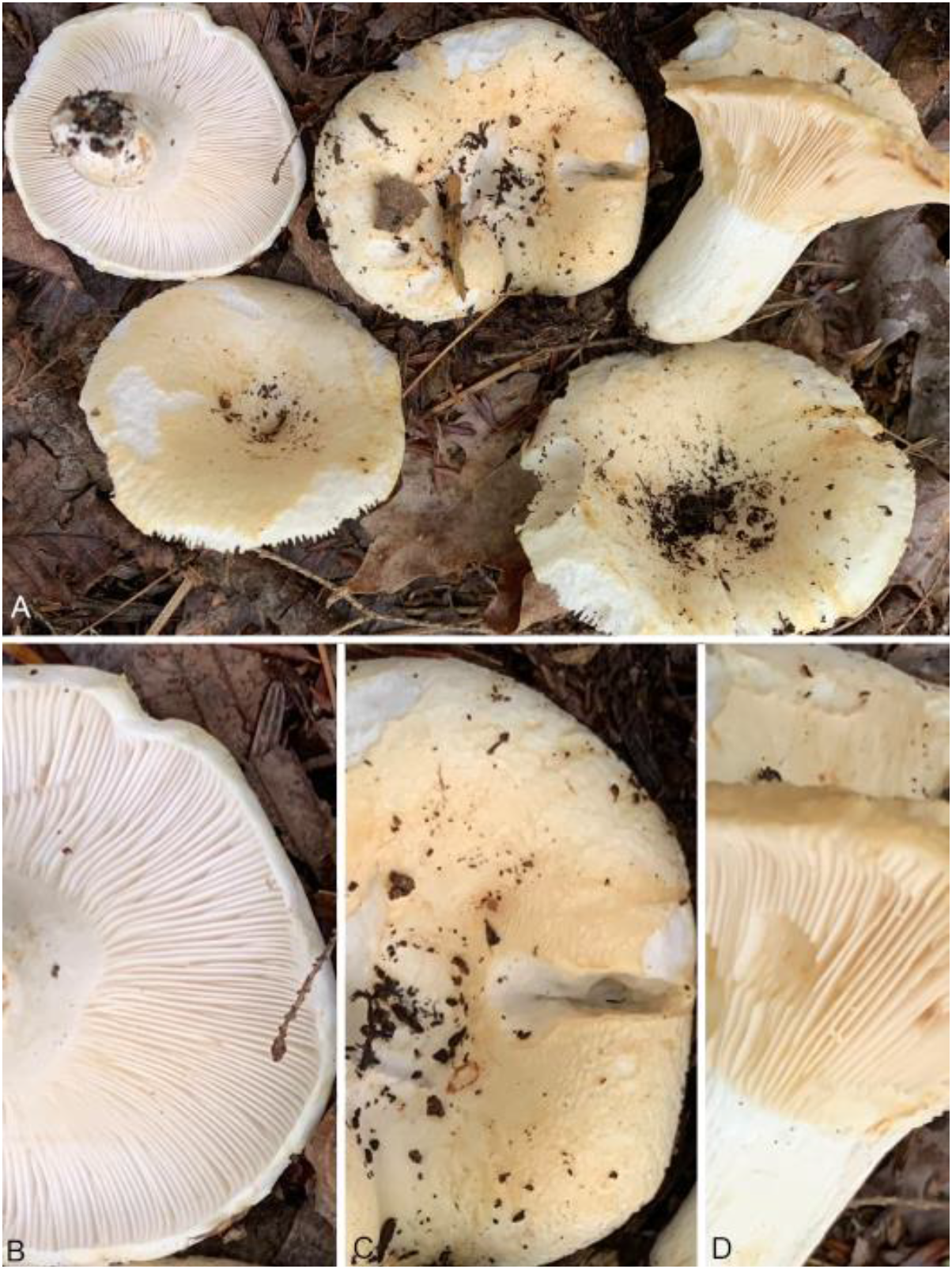
*Russula* cf. *inopina* (*Paula deSanto* MF84640). **A**. General field habit. **B-D**. Details that allow to recognize the species, i.e. almost multiseriate white lamellae when young (**B**), a scurfy pileus margin (**C**) and distinctly yellowish lamellae when mature (**D**). Photo credits P. deSanto.

**Fig. 4.**
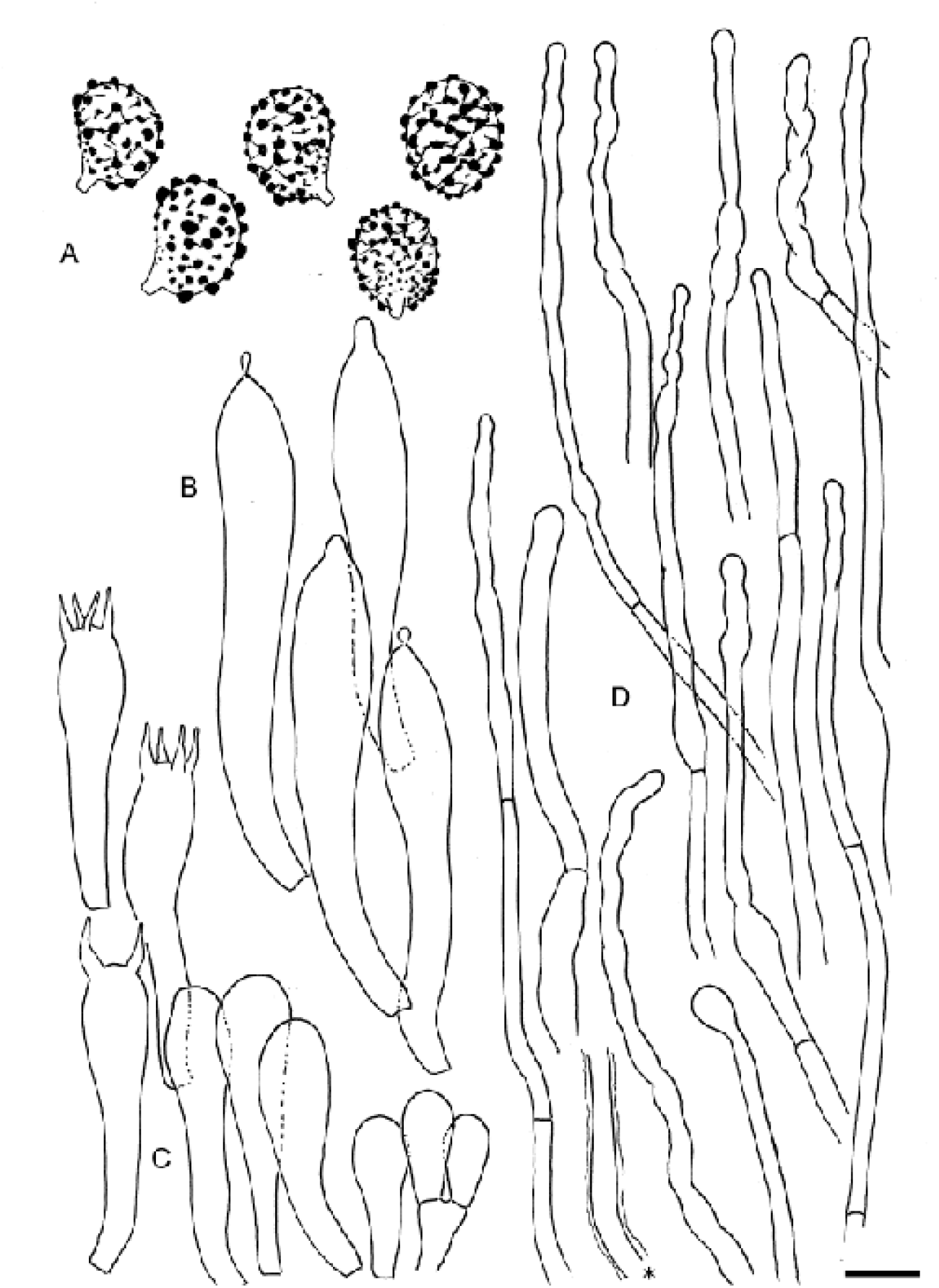
*Russula* cf. *inopina* (*Paula deSanto* MF84640). Microscopic features. **A**. Spores. **B**. Hymenial gloeocystidia that appear optically empty. **C**. Basidia and basidiola. **D**. Hyphal terminations in the pileipellis with * showing a detail of the glutinous sheath observed on many terminations. Scale bar: 10 µm for all elements, except for spores where it is 5 µm. Drawings B. Buyck.

**Spores** (6.2) 6.5−***6*.*9***−7.2 (7.5) × (4.8) 4.9−***5*.*2***−5. (5.6) **µm**, Q = (1.20) 1.27−***1*.*34***−1.40 (1.48), ellipsoid, ornamentation in young spores forming a dense low network, with age developing large hemispherical warts to almost drop-like deposits, mixed with smaller warts of variable sizes and with a variable number of low connections, some spores being almost devoid of connections, others being almost subreticulate; suprahilar spot never distinctly amyloid although sometimes weakly grayish in the distal part, generally finely warty over its entire surface. **Basidia** 35−43 × 8−9 µm, 4-spored, occasionally two-spored near the gill edge, with stout, slender sterigmata, measuring up to 8 × 2 µm. **Basidiola** distinctly clavate and quickly slender. **Hymenial gloeocystidia** 55−80 × 9−10 µm, discrete, originating deeply rooted in the subhymenium, not or hardly emergent, fusiform to narrowly pedicellate-clavate, variably narrowing at apex, sometimes more or less abruptly and almost mamillate at the very apex, others minutely mucronate and droplet-like or minutely globose, more or less thin-walled, without distinct crystalline or granular contents, some distinctly oily-refringent at the apex, negative to sulfovanillin. **Marginal cells** not differentiated, the entire lamellar edges being occupied by basidiola and occasional basidia or cystidia. **Lamellar trama** almost entirely made of large sphaerocytes, near the lamellar edges with dispersed, distinct oleiferous hyphae. **Pileipellis** not metachromatic in Cresyl blue, not clearly two-layered and not distinctly separated from the underlying trama by a much denser tissue (thus indicating it is not separable when fresh), formed of loosely interwoven hyphae, hyphae in the subpellis often clearly incrusted in a typical zebroid pattern, sometimes with distinct mucous sheath. At the pileus surface with long, slender and very narrow, sparsely septate hyphal terminations, mostly only 2−3 µm wide, becoming wider toward the subpellis, many distinctly yellow, cell walls never or only slightly thick-walled; terminal cells (30)50−80 × 2−3 µm, often narrowly cylindrical, with local, slightly and sometimes repeatedly inflated portions, up to 5 µm wide, and very frequently also with slightly to distinctly inflated to nearly globose apex. Pileocystidia doubtful, cells with typical crystalline contents not observed, terminal cells of cystidioid shape because of their minutely mucronate tip rare, but present. Untypical oleiferous hyphae with yellowish but granular, rather than oily contents dispersed in the context. **Clamp connections** absent.

##### Material examined

**USA:** New York, Oneida Co., near Camden, in mixed forest under *Tsuga, Fagus* and *Acer, leg. Paula deSanto MF84640* (CUP 070958; duplicate at PC 0714859).

##### Notes

In the field, the general habit of this American specimen MF84640 immediately suggests affinities with species of *Russula* subgen. *Brevipedum*: firm and fleshy fruiting bodies, short stipe, narrowly depressed center of pileus, and regularly unequal, white lamellae that turn yellowish with age. The distinct yellowish tint of the mature lamellae suggests a coloured spore print, a typical feature of subsect. *Pallidosporinae* of the subgenus. The latter subsection groups all species of subgen. *Brevipedum* with smaller spores, non-amyloid suprahilar spot and a subreticulate, low spore ornamentation composed of locally interconnected, obtuse warts, very different from species in the only other subsection in the subgenus, subsect. *Lactarioideae*. Because of the features of the spores, we could reduce the range of possible candidate species in North America to *R. inopina, R. cascadensis, R. vesicatoria* and *R. angustispora* (Buyck & Adamčík 2013), all known to associate with conifers (Shaffer 1964). *Russula cascadensis* is a rather common West Coast endemic taxon, whereas *R. vesicatoria* occurs on the East coast and around the Gulf of Mexico and differs considerably in the smooth white pileus surface and extremely crowded lamellae; furthermore it is always found under *Pinus* in deep sandy soils. *Russula angustispora* is a species recognizable by its unusually elongate spores and the apparently highly specific association with *P. virginiana*, a conifer found in the southeastern part of the USA (Bills & Miller 1984); furthermore, it produces a deep ochre spore print.

*Russula romagnesiana* Shaffer (Shaffer 1964; Buyck & Adamčík 2013) is another ill-known species in subgen. *Brevipedum* that shares with our specimen a similar spore ornamentation and the same type of hymenial cystidia. It differs, however, in the permanently white lamellae, the amyloid suprahilar spot on the spores and the different shape of the hyphal extremities, which all place *R. romagnesiana* in subsect. *Plorantinae*.

This leaves *R. inopina* as the only possible candidate, unless our collection represents a new species. Because there are some microscopic differences between the type specimen and our collection, we decided to use “*R*. cf. *inopina*” for our specimen. The original description was merely based on two specimens, both collected in early August at close distance near the shores of Burt Lake, Michigan. Shaffer’s description stresses key features shared with our collection, such as the aspect of the pileus margin [“*finely felted on the margin; at times becoming areolately cracked, with the cuticle not separable and the margin not striate”*] and also the very narrow hyphal endings in the pileipellis and the fact that there are irregularly inflated or have globose apices [*“hyaline hairs 2.0-3.0 µm broad and up to 80 µm long which are capitate or taper to blunt apices*”]. Shaffer’s spore measurements are incorrect as we showed in our type study (Buyck & Adamčík, 2013). The correct size of the spores for the type collection is (6.5) 6.9−***7.2***−7.5 (7.8) × (5.0) 5.2−***5.5***−5.7 (6.0) µm, Q=(1.22) 1.25−***1.31***−1.37 (1.44) µm, which matches our observations on the present collection; also the spore ornamentation is very similar although our collection is characterized by the more pronounced development of large, droplet-like ‘warts’, a phenomenon that is frequently observed in occasional specimens of several species belonging to the *Pallidosporinae* or subgen. *Heterophyllidiae*. Even many species of *Lactarius* subgen. *Plinthogali* can show such erratic development. The *R. inopina* holotype differs principally from our specimen in the long basidia and obtuse-rounded apices of hymenial gloeocystidia, two characters that are in our opinion not taxonomically reliable for species recognition.

Another unusual microscopic feature of our specimen is the very poor differentiation of contents in any type of cystidioid cells. This is even true for hymenial cystidia, which are, nevertheless, readily recognized by their different shape as compared to the basidia and basidiola. However, in the pileipellis, morphology is not very helpful in differentiating normal terminal cells from pileocystidia as nearly all hyphal extremities on the pileus surface have more or less granular-refringent or mucilageous deposits or incrustations on their walls. The poor contents of cystidioid elements in our *R*. cf. *inopina* specimen contrasts strongly with the abundant cystidioid contents observed in species of subgen. *Brevipedum*, and even with those observed in the type specimen of *R. inopina* where contents were far from abundant, but nevertheless distinctly present.

The Chinese specimen QC376 grouped with the American *R*. cf. *inopina* and a Japanese environmental sample (GenBank accession LC315898) with ML-BS 71% (Fig. 2). The genetic diversification suggests a different species but since this collection is based on a single fruiting body, we wait for new collections to formally describe it. It is highly similar in morphology to *R*. cf. *inopina*. It has nearly identical sizes for spores [(6.0) 6.5−***6.6***−7.0 × 4.5−***5.1***−5.5 (6.0) µm, Q = (1.08) 1.13−***1.30***−1.40], basidia (37−43 × 7−9 µm) and hymenial gloeocystidia (55−90 × 7−9 µm), the latter being also optically empty or with very sparse granular contents. Although the pileipellis of this Chinese collection did not revive well, we observed undulating hyphal extremities and we can confirm the absence of pileocystidia with distinct contents; also its general field habit (a pale-coloured fruiting body with unequal lamellae) fits the morphological criterion of subgen. *Cremeoochraceae*.

### 2. Russula cremeoochracea R.F.R. McNabb, New Zealand J. Bot. 11(4): 683 (1973)

#### a. *Russula cremeoochracea* var. *cremeoochracea*, Fig. 5

*Type*: **New Zealand**, Prov. Nelson, Oparara, Oparara river, Fenian Track, on soil under *Nothofagus [Lophozonia] menziesii*, 9 Jan. 1968, *leg. R.F.R McNabb* & *E. Horak* 122 (PDD 30661, holotype; ZT 69-309, isotype).

**Fig. 5.**
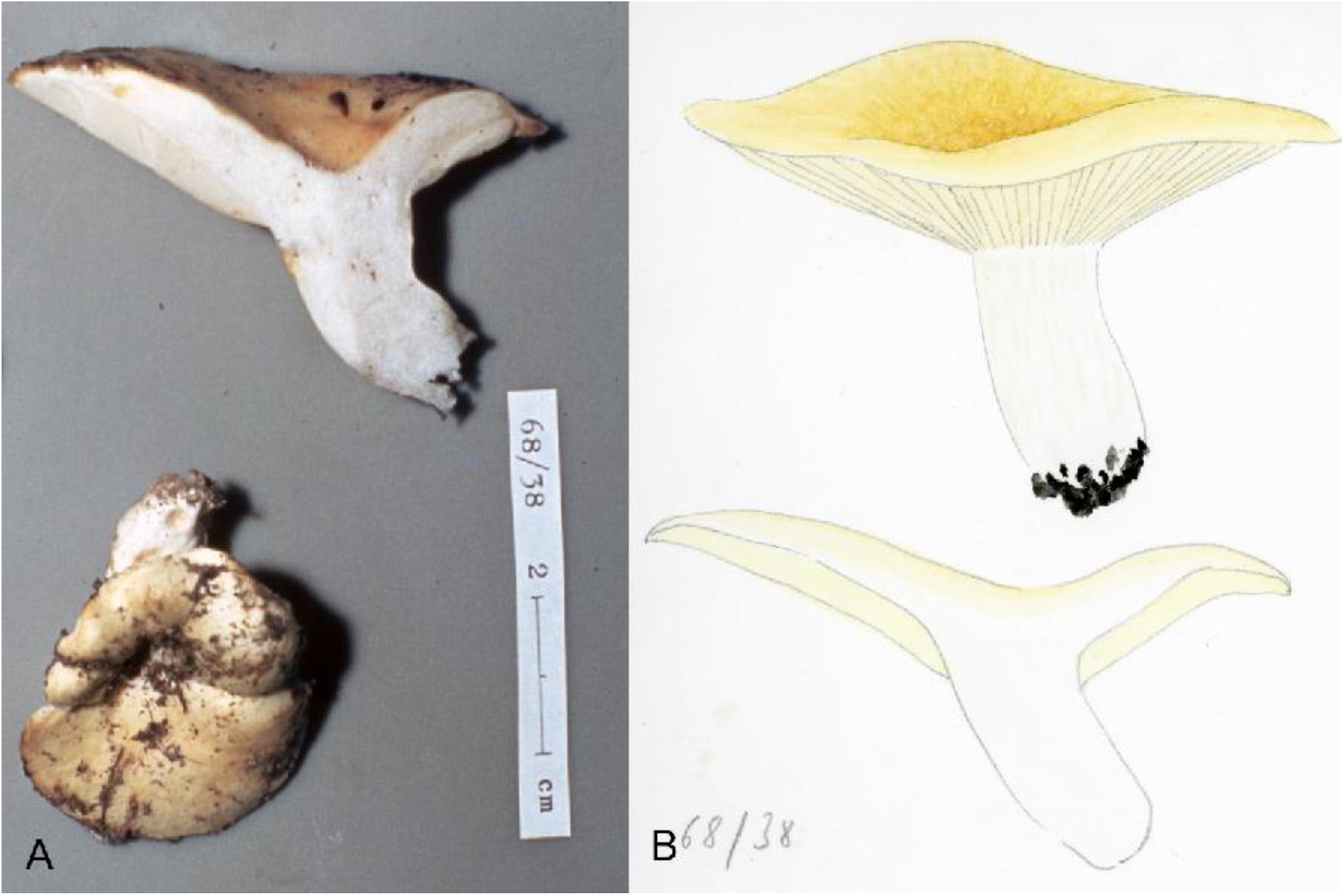
*Russula cremeoochracea* (*E. Horak*, ZT 68-038). Photo credit: E. Horak.

*Other material examined*: **New Zealand**, Prov. Buller, Ngahere, Nelson Creek, 18 Jan. 1968, *leg. E. Horak* (ZT 68-038, with painting and photograph); Prov. Nelson, Lake Rotoiti, Peninsula, under *Nothofagus [Lophozonia]menziesii, Nothofagus [Fuscospora] solandri* var. *cliffortioides, Leptospermum scoparium*, 20 Jan. 1968, *leg. E. Horak* (ZT 69-002).

*Description:* **Pileus** 40−82 mm diam., fleshy, up to 9 mm thick above gill-stipe attachment, 7 mm thick at mid-radius, gently to narrowly depressed in center, involute margin smooth; surface at first cream-colored, becoming butter-yellow or ochraceous (colour like *R. foetens*), dull, dry, slightly sticky when moist, smooth. **Lamellae** irregularly unequal with mostly 1 lamellula/lamella, normally spaced (ca 1/mm), 3−5 mm high, at first white then becoming cream-colored; gill edges entire, concolorous. **Stipe** 25−35 × 16−22 mm, cylindrical or obclavate, smooth, white, compact. **Context** brittle, white, unchanging on exposure. **Odor** not distinctive, weakly of fruit or of camembert cheese. **Taste** mild. **Spore print** pale cream.

**Spores** (5.8) 6.2−***6.5***−6.8 (7.1) × (4.8) 4.9−***5.1***−5.3 (5.4) µm, Q = (1.19) 1.21−***1.28***−1.35 (1.48); ornamentation subreticulate, although often poorly so, in immature spores forming a faintly amyloid, low network, with age developing small hemispherical warts, locally confluent in low crests or more rarely interconnected by low linear tracts, many warts remaining isolated; suprahilar spot inamyloid, although sometimes weakly grayish in the distal part. **Basidia** 36−52 × 8−10 µm, slender, pedicellate-clavate, with (1) 4 slender sterigmata, 5−8 × 1−2 µm, when only 1 or 2 sometimes longer (up to 13 µm) and deformed. **Hymenial gloeocystidia** dispersed, large, 53−114 × 10−13 (17) µm, originating from deep in the subhymenium, highly emergent, spindle-shaped, thin-walled, mostly appendiculate, with indistinct contents that have the aspect of a cloudy, oily-refringent mass that is glued to the wall but leaving dispersed circular “holes”. **Lamellar trama** composed of sphaerocytes that rarely are circular and form a dense tissue, intermixed with some hyphae; oleiferous fragments present. **Subhymenium** densely pseudoparenchymatic. **Lamellar edge** occupied by basdiola and cheilocystidia. **Pileipellis** not metachromatic in Cresyl blue, not clearly two-layered and not abruptly separated from the underlying trama by a much denser tissue (thus indicating it is not separable when fresh), composed of loosely interwoven hyphae, in the subpellis with the larger cells often obscurely incrusted in typical zebroid pattern, sometimes with distinct mucous sheath; suprapellis formed of long, multi-celled, branching hyphal terminations on subcylindrical cells, 4−8 µm wide; the branching cells often variously inflated (sausage-like, clavate, barrel-shaped, ellipsoid, utriform or vesicle-like, etc.) and up to 15 µm diam.; terminal cells mostly (10) 30−50 µm long, usually slightly narrowing, often undulate-wavy in outline, obtuse to subcapitate at apex. Toward the pileus center with much longer (up to 100 µm) terminal cells that are only 2−3 µm wide, with subcapitate, moniliform to even inflated apex. Pileocystidia not abundant, one-celled, thin-walled, usually with ill-differentiated contents or optically empty, SV-negative, recognizable by their needle-shaped to conical shape and minute appendiculate to moniliform or mucronate apices. Untypical oleiferous hyphae with yellowish but granular-oily content dispersed but rare in context. **Clamp connections** absent.

*Notes*: The description of the microscopic features is mainly based on the isotype, which consists of a very small portion of a pileus from the holotype collection. The features of the isotype agree perfectly with those of both other collections studied here. McNabb’s species resembles in many ways *R. inopina* under the microscope because of the similar aspect of hyphal extremities in the pileus and the similar size and ornamentation of the spores. *Russula cremeoochracea* has always been described as lacking pileo- and caulocystidia (McNabb 1973; Cooper & Leonard 2014). However, dermatocystidia are actually present, both on the pileus and on the stipe surface, but are indeed difficult to trace because many are optically empty or have poorly visible contents that do not react in SV. In all examined collections, however, these pileocystidia are readily recognized because of their upwards attenuating shape ending in a minutely capitate to moniliform apex.

*Russula cremeoochracea* shares with our specimen of *R*. cf. *inopina* also the often obclavate stipe with broadly rounded base, which is not rooting as in some species of subgenera *Compactae* or *Archaeae* (see for ex. Buyck *et al*. 2018, fig. 7a). The white lamellae turning yellowish with age is a feature that is more pronounced in *R*. cf. *inopina* and the latter also has much more lamellulae compared to McNabb’s species.

Contrary to the extremely rare American *R. inopina, R. cremeoochracea* seems to be widely distributed in New Zealand as suggested by more than 40 collections reported to be kept in Herbarium PDD in Auckland (https://nzfungi2.landcareresearch.co.nz). However, molecular verification might be in order as we will show below that there exist two distinct entities depending on the associated host. McNabb (1973) did not distinguish between these and his description was based on specimens collected both under *Nothofagaceae* and *Myrtaceae*. Furthermore, this species seems easily confused in the field with several others among McNabb’s species, principally with those from the *R. allochroa/australis/multicystidiata* complex in subgen. *Brevipedum*, and with *R. littorea* from subgen. *Crassotunicatae*.

### *Russula cremeoochracea* var. *myrtacearum* Buyck & X.H. Wang, var. nov. Mycobank: xxxxxx. Figs. 6, 8c–g, 9

*Etymology*: named after its host association with trees in *Myrtaceae*

**Fig. 6.**
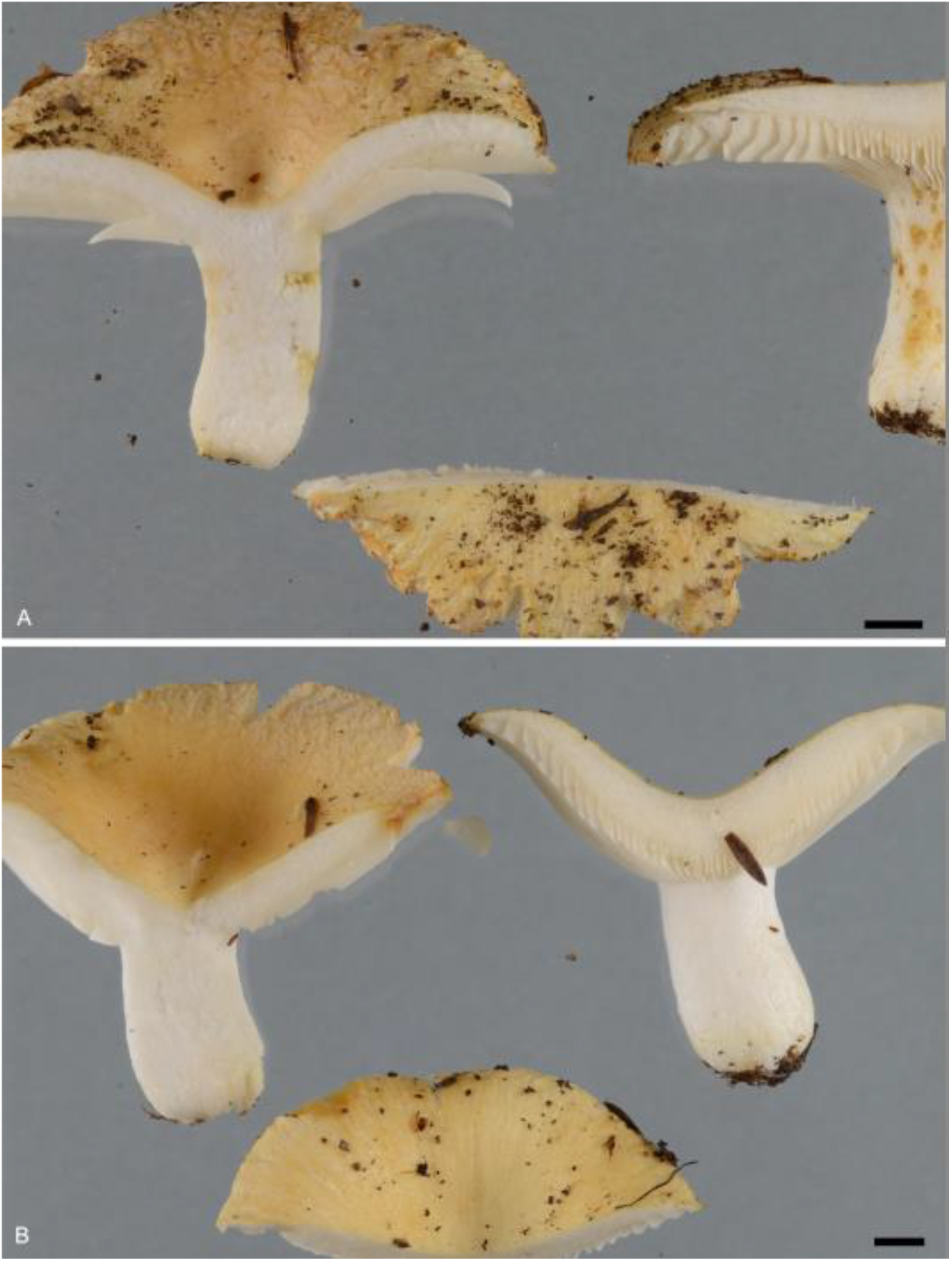
*Russula cremeoochracea* var. ***myrtacearum*, var. nov. A**. PDD104171. **B**. PDD104175. Scale bars: 5 mm. Photo credits J. Cooper

**Fig. 7.**
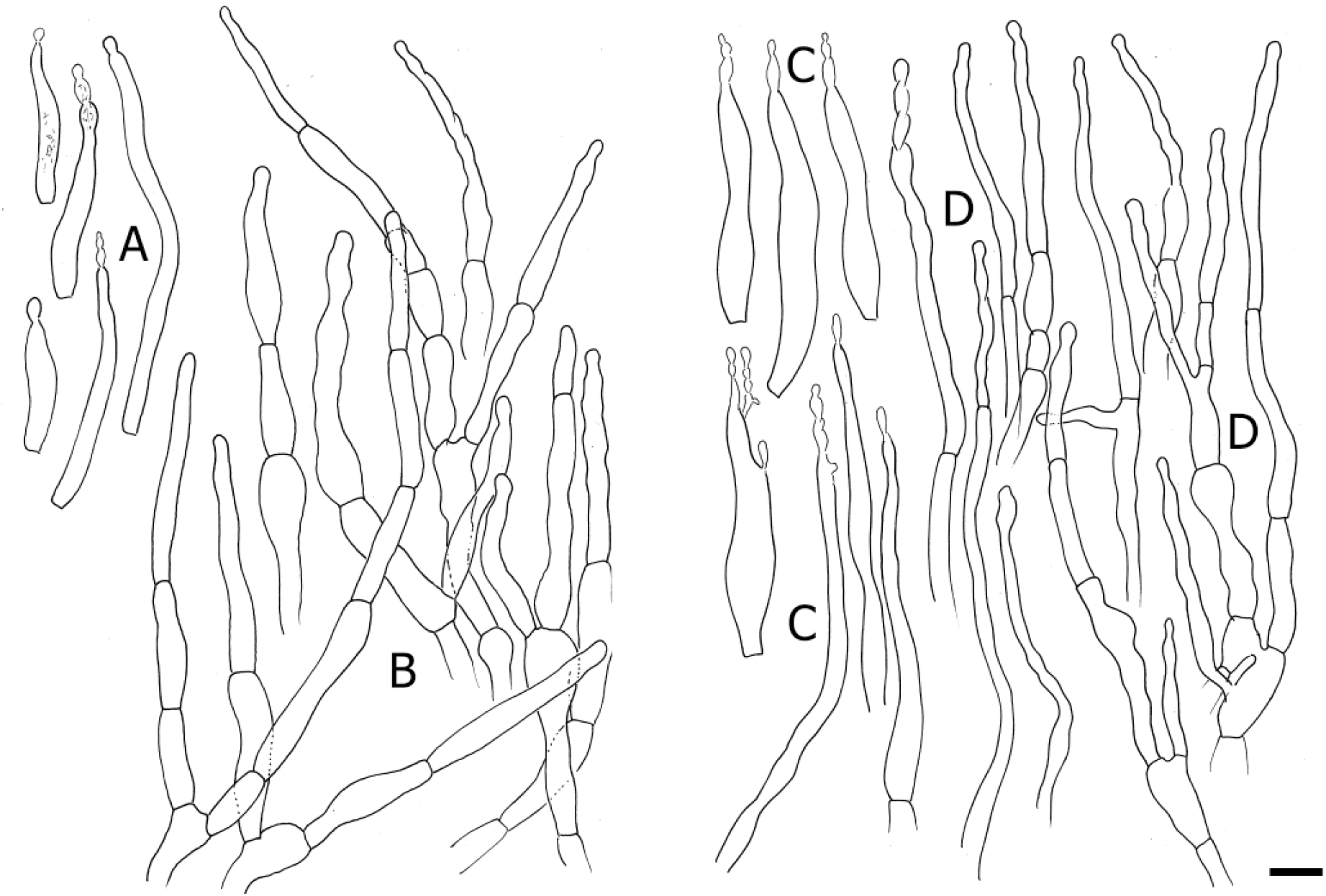
*Russula cremeoochracea* var. *cremeoochracea* (E. Horak, ZT 69-309, isotype). **A**. Pileocystidia with hardly any contents near pileus margin. **B**. Hyphal terminations near pileus margin. **C**. Pileocystidia with hardly any contents in pileus center. **D**. Hyphal terminations in pileus center. Scale bar: 10 µm. Drawings B. Buyck.

**Fig. 8.**
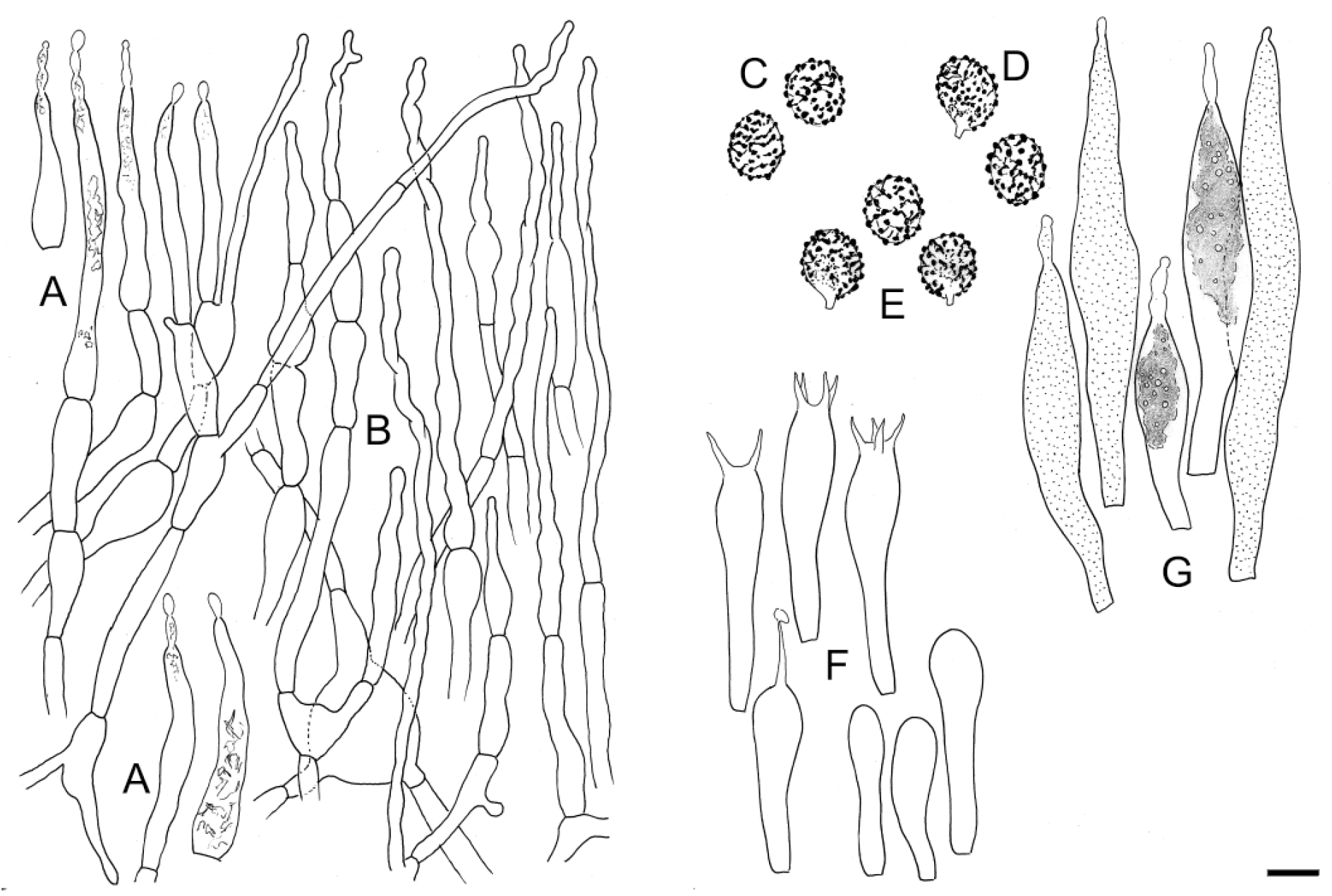
*Russula cremeoochracea* var. *myrtacearum* var. nov. (ZT 68-365). **A**. Pileocystidia. **B**. Hyphal terminations in the pileipellis. **C-E**. Spores of *R. cremeoochracea* var. *cremeoochracea* in Melzer’s reagent (a, ZT 69-309, isotype; b, ZT 68-038); **F**. Basidia and basidiola. **G**. Hymenial cystidia. Scale bar: 10 µm, but only 5 µm for spores. Drawings B. Buyck.

**Fig. 9.**
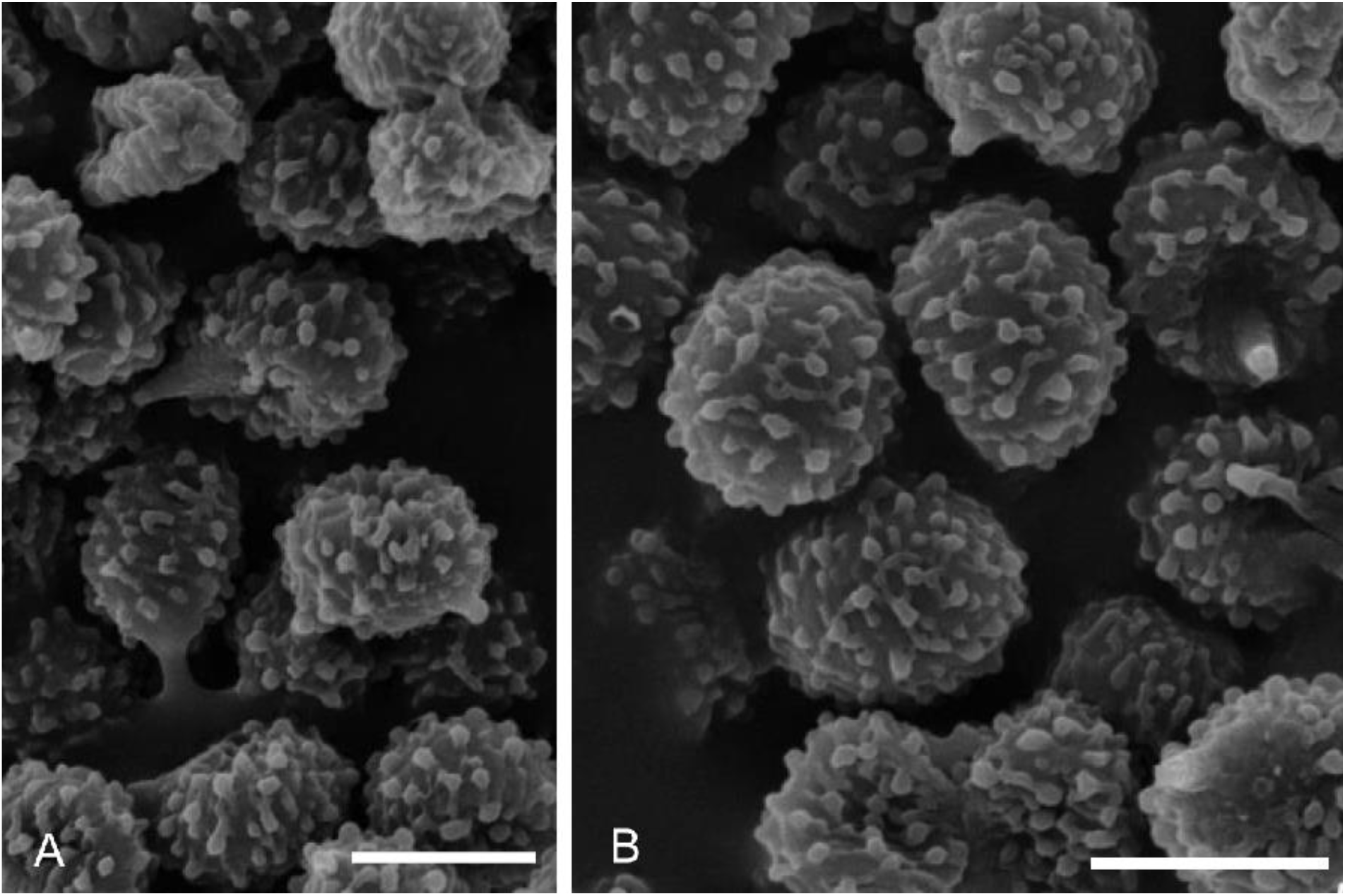
*R. cremeoochracea var. myrtacearum*. Spores under scanning electron microscope. **A**. PDD104171. **B**. PDD104175. Scale bars: 10 µm. Photo credits J. Cooper.

*Diagnosis*: differs from the type variety in the bright naples or mustard yellow (3A6-7) and somewhat less fleshy pileus, which is strongly felty-scurfy, becoming distinctly areolate-cracked toward the margin; furthermore, it associates with *Myrtaceae* instead of *Nothofagaceae*, and has several base pairs differences in both ITS and LSU sequence data.

*Holotype***: New Zealand**, Auckland, Atkinson Park, under *Kunzea (Leptospermum) ericoides* and *Agathis australis*, 31 May 1969, *leg. E. Horak* (ZT 69-365).

*Additional material examined*: **New Zealand**, Auckland, Waitakere Ranges, near start of Parau Track and Farley Track, 2650122 E – 6466446 N (WGS84 -36.993321 174.570115), under *Kunzea ericoides*, 29 Jan. 2014, *leg. P.R. Johnston, J.M. Ryder, O.K. Sigglekow, B.C. Robson* s.n. (PDD104171); idem (PDD104175)

*Description:* **Pileus** 40–50 mm diam., concave with a distinct central depression from the beginning, margin young involute; surface young cream coloured, turning to bright naples or mustard yellow (3A6-7) and distinctly scurfy-felty when mature, becoming areolate-cracked with whitish patches or even radially fissuring toward the margin, dry but viscous when humid. **Lamellae** adnate, unequal from irregularly inserted shorter lamellulae, 2–3 mm high, white when young, turning to cream with age; edge concolorous, even. **Stipe** 16–18 × 10–13 mm, shorter than the pileus diameter, cylindrical or slightly widening toward the base, smooth, ivory white, not lacunar inside. **Context** white, brittle, not changing colour with age, but turning yellowish to rusty brown where injured. **Odor** none or faintly reminiscent of camembert cheese. **Taste** mild. **Spore print** not obtained.

### Spores (6.0) 6.2−*6.5*−6.8 (7.1) × (4.6) 4.8–*5.0*–5.3 (5.7) µm, Q = (1.19) 1.23−*1.30*−1.36 (1.42), ellipsoid. Other features as in the type variety

*Notes*: McNabb’s original description of *R. cremeoochracea* describes a smooth pileus surface [“glabrous, faintly innately pruinose under lens, velar remnants absent”], a feature that is confirmed by the notes taken by E. Horak on the same type collection. In contrast to these features of the type variety, the strongly felty-scurfy, irregularly cracked pileus surface of this new variety is unmistakable (Fig. 6) and it is also of a more intense yellow colour. Microscopic features, however, appear to be near-identical.

The two varieties differ by five and four consistent differences in their ITS and LSU sequences respectively resulting in a 99% similarity between them. There are also differences in the protein-coding genes and mtSSU region, but there is only a single specimen sequenced for the var. *myrtacearum*. The two varieties might argue in favour of the recognition of two distinct species, each associated with a different host tree family. However, we think it is necessary to collect and study more specimens before this decision can be made.

## DISCUSSION

### Strange BLASTn results for ITS suggest ancient or isolated lineages

BLASTn of ITS sequences for species in ancient lineages can be quite disorientating. We already discussed this aspect when describing the new subgen. *Glutinosae* (Buyck et al. 2020). In that subgenus, the first 100 BLAST hits for nrITS sequences (arranged by max score and excluding environmental sequences) did not list a single species that belongs to either *Compactae* or *Archaeae*, which constitute the two closest subgenera to subgen. *Glutinosae* (supported by very high bootstrap values: ML BS=99%); on the contrary, all of the other subgenera showed up in the first 100 BLASTn results. In the case of *R*. cf. *inopina*, the biggest surprise came from BLASTn of the obtained ITS sequence. In contrast with its high morphological similarity to typical species of subgen. *Brevipedum*, BLASTn results were exclusively on subsect. *Ochroleucinae* (*Russula ochroleuca, R. viscida, R. vinacea, R. krombholzii, R. pumila*) belonging to the *Russula* subgen. *Russula* core clade (85% similarity for 100% coverage), with further hits on subsect. *Subvelatae* (*Russula insignis, R. pulverulenta*) from subgen. *Heterophyllidiae* sect. *Ingratae*. Nothing was more similar in the first 250 BLASTn results when sorted according to e-value. When sorting the BLASTn results by ‘percentage identity’, the closest hit was then on *R. pulverulenta* (89.29% for 71% coverage) but still not a single species from subgen. *Brevipedum*, except for a *R. afrodelica* from Madagascar (87.57% for 87% coverage), a species that is otherwise completely different from a micromorphological point of view. The first species (*R. pallidospora*) from subgen. *Brevipedum* shows up in ITS BLASTn results below 84% similarity together with species from other subgenera. The fact that our multi-locus phylogeny placed the new subgenus in superclade II might explain this strange BLASTn result as relations between most of the subgenera within this superclade remain unresolved.

When blasting against environmental sequences, there are two additional close hits: the first (LC315898, 95.28% similarity) was obtained from an ectomycorrhizal sample associated with the highly endangered endemic *Pinus amamiana* in Japan (Murata *et al*. 2017); the second closest hit (GQ268654, 92.24% similarity) corresponded to an ectomycorrhizal root tip from dipterocarp forest in Sarawak, Borneo (Peay *et al*. 2010). Finally, in 2021, two ITS sequences were deposited for documented collections of *R. cremeoochracea* from New Zealand (see Fig 6). These sequences now constitute the second closest hit with 93.11% similarity for 100% coverage with *R*. cf. *inopina*, while no other sequences are more than 86.6% similar.

### Towards a more precise image of the ancestral *Russula*

With the present description of a new subgenus *Cremeoochraceae*, the number of subgenera in *Russula* amounts now to nine, seven of which are characterized by unequal gills or harbour at least some species with unequal gills. The presence of unequal gills is a feature that in traditional European monographs opposed subgen. *Compactae* to the rest of the genus. Regularly polydymous gills, which are typical of *Lactarius* and *Lactifluus*, are found in subgenera *Compactae, Glutinosae* and *Brevipedum*, and also characterize some species in *Malodorae* and now *Cremeoochraceae*. Irregularly unequal gills are typical for subgenera *Archaeae* and *Crassotunicatae*, but characterize also the type species in *Cremeoochraceae*, as well as isolated species in *Heterophyllideae* (e.g. subsection *Aureotactinae*) and even in the core clade of subgen. *Russula* (e.g. some rare species in subsect. *Emeticinae*). Almost regularly forked gills, so typical for all *Multifurca* (Wang *et al*. 2018) and also observed in some *Lactifluus*, are present in some species of subgen. *Malodorae* (e.g. sect. *Edules*) and in some *Heterophyllidiae* (e.g. some species of subsect. *Cyanoxanthinae*). The occurrence of equal gills is restricted in the entire family Russulaceae to the overwhelming majority of the *Russula* species in the two most speciose subgenera, *Russula* and *Heterophyllidiae*.

Spore print colour is another intriguing feature in the family. White spore prints have always been interpreted in the more traditional monographs of the genus (Romagnesi 1967, Sarnari 1998, 2003) as representative of the most ancient condition, not only in the genus, but even in the entire family. In recent years, the very dark spore prints that characterize all species in *Multifurca* (Wang *et al*. 2018) and the coloured spore prints of the two most recently described ancient subgenera in *Russula*, viz. *Glutinosae* and now also *Cremeoochraceae*, question this hypothesis. Also in *Russula* subgen. *Brevipedum* the most basal lineages are those with coloured spore prints. All of the species in the other subgenera in *Russula* produce either white to whitish spore prints (*Archaeae, Compactae, Crassotunicatae, Malodorae*) or have developed more coloured spore prints in terminal clades (*Heterophyllidiae, Russula*). The ancestral state reconstruction based on a three-locus phylogeny (ITS, mtSSU, *rpb2*) published by Cabon *et al*. (2017) suggested a dark yellow spore print for the ancestor of the *Russula* subgen. *Russula* crown clade. This, however, was clearly a premature conclusion based on an insufficient representation of existing lineages. Buyck *et al*. (2018) demonstrated in their five-locus phylogeny that the four most basal, known lineages in subgen. *Russula* crown clade are composed of pale-spored species. Also the subgen. *Russula* core clade is mainly composed of pale-spored lineages.

With very few exceptions, fruiting bodies formed by species in subgenera *Archaeae, Glutinosae, Cremeoochraceae* and *Brevipedum* are pale-coloured (white to yellowish). In subgen. *Compactae*, whitish or very pale colours are typical in early stages of many species or stay present until maturity as, for example, in the type species of its sect. *Polyphyllae*. Bright red, orange, blue and green colours are restricted to more terminal clades in subgenera *Russula* and *Heterophyllidiae*, which implies that these colours are definitely acquired in more terminal clades.

Host association is a feature that remains difficult to interpret. Species belonging to our new subgenus definitely associate with both coniferous (the environmental sample under *Pinus* in Japan) and deciduous trees (the environmental sample from dipterocarp forest in Borneo, as well as *Nothofagus* and Myrtaceae in New Zealand).

Based on the above, we can hypothesize that the ancestral *Russula* is an agaricoid, pale coloured species with unequal gills, without suprahilar amyloid spot and low spore ornamentation. Other features need further investigation at this time.

### Subgen. *Cremeoochraceae* exhibits a circum-Pacific distribution pattern and adds support to a non-African origin of the genus

Looney *et al*. (2016) speculated that *Russula* was an ancestrally temperate lineage, but the sampling did not include representatives of *R*. subgen. *Glutinosae* and the new subgenus described here, and did not place subgen. *Crassotunicatae* with significant support. Their ancestral reconstruction was largely based on ITS metadata, thus generating a different topology from Buyck *et al*. (2018, 2020) and our multi-locus phylogenetic analyses (Fig. 1). As we have demonstrated above, using similarity of ITS sequences for phylogenetic inference might be disorientating. Although subgenera *Glutinosae, Crassotunicatae* and *Cremeoochraceae* are each holding less than a handful of species, their phylogenetic and biogeographical implications cannot be ignored. Our new subgenus, as shown in the ITS-LSU tree (Fig. 2), is composed of at least six phylogenetic entities. It is absent from Europe and Africa and together, these five species exhibit a circum-Pacific distribution pattern, although still with the exclusion of South America. A similar distribution pattern is observed in *Multifurca* (Wang *et al*. 2018). Ancestral Area Reconstruction suggested that *Multifurca* mostly likely had its origins in Australasia and this seems also the case for our new subgenus which is predominantly distributed in Australasia with 5 out of 6 taxa.

*Lactifluus*, the sister genus of *Russula*, is overwhelmingly more diverse in the tropics (De Crop *et al*. 2017), and even *Lactarius*, a genus predominantly occurring in temperate or colder climates (Buyck *et al*. 2010), harbours more tropical species than previously thought and these are usually sitting on long branches in phylogenetic analyses (Wang *et al*. 2020; Wu *et al*. 2022). Hackel *et al*. (2022) inferred a tropical African origin for the three major ectomycorrhizal genera in family Russulaceae, as well as a tropical African origin for the large majority of the individual subgenera in *Russula*. The latter conclusion, however, is not at all what is suggested by our multigene analysis. On the contrary, none of the nine subgenera in *Russula* is here suggested to have an African origin. One could argue that this might be the result from limited sampling, but even recent studies with extensive sampling within individual subgenera appear to suggest a similar scenario. Example given, Xie *et al*. (2023) described the tropical Asian *R. lacteocarpa* as basal to all known species in subgen. *Archaeae*, one of the more ancient lineages in the genus. Three of the other most ancient lineages in the genus, subgenera *Glutinosae, Crassotunicatae* and *Cremeoochraceae* are even completely absent from Africa and share an amphipacific distribution pattern with *Multifurca*, the latter being definitely a good candidate for oldest ectomycorrhizal lineage in Russulaceae due to shared morphological similarities with some of the corticioid genera in the family (Buyck *et al*. 2008).

Our phylogeny offers now support to distinguish two main superclades within *Russula*: full ML-BS support for a lineage that contains subgenera *Glutinosae, Archaeae* and *Compactae*, and 80% ML-BS support for the superclade that contains all other subgenera. The former is at the moment definitely not of African origin, whereas the second is equally suggested as not having African origins. The new subgenus *Cremeoochraceae*, unrecorded from Africa, South America and Europe, is peculiar in that it harbours both tropical, warm- and cold-temperate species among the handful of potential taxa it contains. Indeed, ITS sequences (see our Fig. 2) suggest that it is present in tropical Borneo and Malaysia, while it occurs in the northern USA, close to the Canadian border, in warm-temperate Japan and in the temperate *Nothofagus* and Myrtaceae forests of New Zealand. In this respect, our new subgenus differs from two other, very small and ancient subgenera: *Glutinosae* with an exclusively northern hemisphere distribution, and also *Crassotunicatae*, a subgenus that inhabits temperate climates of the northern hemisphere but is also present in New Zealand (*R. littorea*). Broader sampling, especially in Australasia, will continue to contribute to answer the origin of Russulaceae and of *Russula* in particular.

## ACKNOWLEDGEMENTS

This work is supported by the National Natural Science Foundation of China (No. 32170022), the Biodiversity Survey and Assessment Project of the Ministry of Ecology and Environment, China (2019HJ2096001006), the project of Investigation of Macrofungi of Maguan County issued by Ministry of Ecology and Environment of the People’s Republic of China and the CAS Key Laboratory for Plant Diversity and Biogeography of East Asia, Kunming Institute of Botany (KIB), Chinese Academy of Sciences (CAS) (project no. LPB201501). We thank Dr. Q. Cai and Dr. G. Wu, KIB) CAS for the specimens; S.Q. Cao and W.B. Li, KIB, CAS helped with lab work and data analysis. Fungarium of Guangdong Institute of Microbiology (GDGM), China and Dr. H.J. Li, National Institute of Occupational Health and Poison Control, Chinese CDC, Beijing, China generously provided loans of *R. ochrobrunnea* for our study. Paula deSanto from the Central New York Mycological Society and colleagues are thanked for bringing the new specimen of *R*. cf. *inopina* to our attention and providing photographs.

## Notes

### Competing Interest Statement

The authors have declared no competing interest.

